# Unmasking the tissue microecology of ductal carcinoma *in situ* with deep learning

**DOI:** 10.1101/812735

**Authors:** Priya Lakshmi Narayanan, Shan E Ahmed Raza, Allison H. Hall, Jeffrey R. Marks, Lorraine King, Robert B. West, Lucia Hernandez, Mitch Dowsett, Barry Gusterson, Carlo Maley, E. Shelley Hwang, Yinyin Yuan

## Abstract

Despite increasing evidence supporting the clinical relevance of tumour infiltrating lymphocytes (TILs) in invasive breast cancer, TIL spatial distribution pattern surrounding ductal carcinoma *in situ* (DCIS) and its association with progression is not well understood.

To characterize the tissue microecology of DCIS, we designed and tested a new deep learning pipeline, UNMaSk (UNet-IM-Net-SCCNN), for the automated detection and simultaneous segmentation of DCIS ducts. This new method achieved the highest sensitivity and recall over cutting-edge deep learning networks in three patient cohorts, as well as the highest concordance with DCIS identification based on CK5 staining.

Following automated DCIS detection, spatial tessellation centred at each DCIS duct created the boundary in which local ecology can be studied. Single cell identification and classification was performed with an existing deep learning method to map the distribution of TILs. In a dataset comprising grade 2-3 pure DCIS and DCIS adjacent to invasive cancer (adjacent DCIS), we found that pure DCIS cases had more TILs compared to adjacent DCIS. However, TILs co-localise significantly less with DCIS ducts in pure DCIS compared with adjacent DCIS, suggesting a more inflamed tissue ecology local to adjacent DCIS cases.

Our experiments demonstrate that technological developments in deep convolutional neural networks and digital pathology can enable us to automate the identification of DCIS as well as to quantify the spatial relationship with TILs, providing a new way to study immune response and identify new markers of progression, thereby improving clinical management.

## Introduction

Ductal carcinoma *in situ* (DCIS) is a non-obligatory precursor of invasive breast cancer (IDC). It is the most common mammographically detected breast cancer, however, predicting DCIS progression to IDC remains a major clinical challenge (1-3). A recent study based on evolutionary models has categorized DCIS evolution to IDC into four models, highlighting its heterogeneity (3). Histology slides enable pathologists to evaluate complex tissue growth patterns in terms of tissue morphology and architecture to arrive at a diagnosis. DCIS lesions are composed of malignant epithelial cells which proliferate within the breast terminal duct lobular unit and are surrounded by myoepithelial cells and basement membrane. The architectural pattern of DCIS is highly variable and is broadly comprised of solid, cribiform, papillary and comedo type of DCIS (4). Such diverse patterns of DCIS present challenges for achieving good inter-observer as well as intra-observer reproducibility in diagnosing and discriminating prognostic features of DCIS (5). Machine learning tools to supplement the current pathology review process could improve objective histologic assessment of DCIS and allow for more reproducible discrimination of good, from poor prognosis in DCIS.

Deep learning has emerged in artificial intelligence to bring powerful tools to many applications, including digital pathology (6-8). To the best of our knowledge, only few automated methods based on machine learning have been proposed for evaluation of DCIS on haematoxylin and eosin (H&E) samples. One of the approaches used multiscale superpixels to discriminate epithelial area from the remaining tissue area and further clustered the epithelial regions based on a random polygon model, but had difficulty with comedo DCIS (9). More recently a new deep learning pipeline was developed to predict risk of recurrence in pure DCIS patients treated with breast-conserving surgery, where texture features were utilised for classification of image patches into normal duct, stroma, cancer duct and immune rich patterns in H&E (10). However, such efforts have not yet been applied in the studies of DCIS to evaluate the spatial variability related to individual DCIS ducts with single-cell spatial resolution, or the relationship of these ducts to the immune microenvironment.

Recent studies support the importance of tumour infiltrating lymphocytes (TILs) in the progression from DCIS to IDC (11) and risk of local and metastatic recurrences (12). However, TIL assessment still relies heavily on quantification by pathological scoring (13), which is labour intensive and often averages over the entire slide. DCIS often involves multiple ductules within a single lobule. When this occurs multiple ductules are often seen as individual duct like structures filled with cancer cells and surrounded by a myoepithelial layer and basement membrane, but separated by connective tissue (14). In this study we aimed to study individual ductules and their surrounding lymphocytes and have adopted the terminology of DCIS duct for each individual ductule examined. Given the complex spatial ductule structure, ecological dynamics between individual DCIS ducts and their surrounding TILs are difficult to measure by eye. These ultimately limit our ability to study the interplay of these cells within the local microenvironment and its impact on tumour evolution and response to treatment (15).

To enable spatial mapping of TIL distribution patterns for individual DCIS ducts in H&E, we designed an integrated computational framework using deep learning. We hypothesise, and provide preliminary data, that the local micro ecology for individual DCIS within the tissue creates differential selective forces and may ultimately influence its potential for progression to invasive cancers. Our primary aims were: 1) to develop and validate a computational pipeline that accurately detects and segments individual DCIS ducts; 2) to characterize the local micro ecology for each DCIS duct, 3) to test the difference in DCIS microecology between samples with pure DCIS disease and samples derived from IDC patients (adjacent DCIS).

## Materials and Methods

### Datasets

We utilise three independent breast tumour datasets which we refer to as the Duke, TransATAC and IHC datasets. These samples were processed, stained and scanned in independent laboratories using different digital slide scanners.

#### Duke Dataset

The Duke dataset consists of samples of pure DCIS disease and DCIS with adjacent invasive cancer, the latter serving as an indicator for poor prognosis DCIS based on the fact that progression to invasive cancer has already occured. H&E stained images acquired from subjects with no invasive component are henceforth referred as pure DCIS and images from patients with an invasive component along with DCIS are henceforth referred to as adjacent DCIS throughout the manuscript. 140 WSIs were obtained from formalin-fixed paraffin-embedded blocks from 65 patients (n = 40 pure and n = 25 adjacent), ranging from 1-3 slides per patient. These were digitized with an automated whole-slide Aperio scanner at a resolution of 0.5 µm/pixel at 20x magnification. The study was approved by the institutional review board of Duke with a waiver of the requirement to obtain informed consent. Patient demographics and baseline characteristics of the Duke dataset are summarized in Table 1.

**Table 1.** Demographics of patients belonging to DUKE dataset comprising pure DCIS and Adjacent DCIS.

| Features | Pure DCIS | Adjacent DCIS |
| --- | --- | --- |
| **Age (mean ± std)** | 56.19 ± 11.47 | 61.26 ± 7.40 |
| >60 | 18 | 14 |
| <60 | 22 | 11 |
| **DCIS Grade** |  |  |
| 2 | 17 | 7 |
| 3 | 23 | 17 |
| 2-3 |  | 1 |
| **ER Status** |  |  |
| Positive | 27 | 16 |
| Negative | 7 | 9 |
| Not Processed (NP) | 6 | 0 |
| **PR Status** |  |  |
| Positive | 27 | 14 |
| Negative | 7 | 11 |
| Not Processed | 6 | 0 |
| **HER2 Status** |  |  |
| Positive | NA | 5 |
| Negative | NA | 20 |

#### TransATAC Dataset

TransATAC is the translational clinical study of the Arimidex, Tamoxifen, Alone or in Combination (ATAC) trial on postmenopausal patients with ER+ breast cancer treated with tamoxifen or anastrozole (16). 30 WSI H&E images representing 30 invasive breast carcinomas in the TransATAC study were selected which contained adjacent DCIS. The study was approved by the South-East London Research Ethics Committee, and all patients included gave informed consent. The images were digitally scanned using Hamamatsu Nanozoomer scanner at a pixel resolution of 0.45µm/pixel at 20x magnification.

#### IHC Dataset

A set of 8 samples were inspected by pathologists and selected on the basis that they contained confirmed areas of DCIS and IDC in the same tissue specimen as part of a previous study (17). H&E staining and immunohistochemistry using CK5 were performed on consecutive 3µm thick sections of paraffin embedded blocks and digitalised with Hamamatsu Nanozoomer whole slide scanner at a pixel resolution of 0.45µm/pixel at 20x magnification.

### Manual annotation collection protocol

Ground truth annotations for DCIS ducts were hand marked by an expert pathologist (B.G.) on the WSI acquired from both Aperio and Hamamatsu scanners. We then used customized scripts to export raw annotations from Aperio Imagescope. These annotations of DCIS, together with annotations of single cells (epithelial cells, lymphocytes, fibroblasts), were used for training deep learning networks. Subsequently, we split each of these WSI image into tiles of size 2000×2000 pixels and the respective masks of the corresponding tiles were used in our pre-processing. Quality control of annotations were performed at the tile level and expert consensus was obtained before using the respective tiles for training. The tiles were then made into patches of different size to cater to our training network.

### Training and validation of DCIS deep learning

We randomly selected 20 WSI each from the Duke and TransATAC datasets for training the deep learning methods. The validation dataset consisted of 10 WSI from Duke and 10 WSI from TransATAC. 10 WSI with annotation from Duke was used as test dataset. Furthermore, 100 and 8 WSI were held out as independent test WSIs from the Duke and the IHC dataset, respectively. Further breakdown of number of annotated tiles employed in training, validation and testing are tabulated in Table 2. Representative images from these datasets indicating varying growth patterns and DCIS surrounded by different microenvironments are shown in Table 3. Training images include both positive and negative examples. Negative examples were from the adjacent DCIS comprising invasive regions, pure stromal and background regions. Balanced number of samples based on presence and absence of the mask were estimated automatically and sampled based on lesser number belonging to the positive image to circumvent the data imbalance between the positive and negative representation.

**Table 2.**
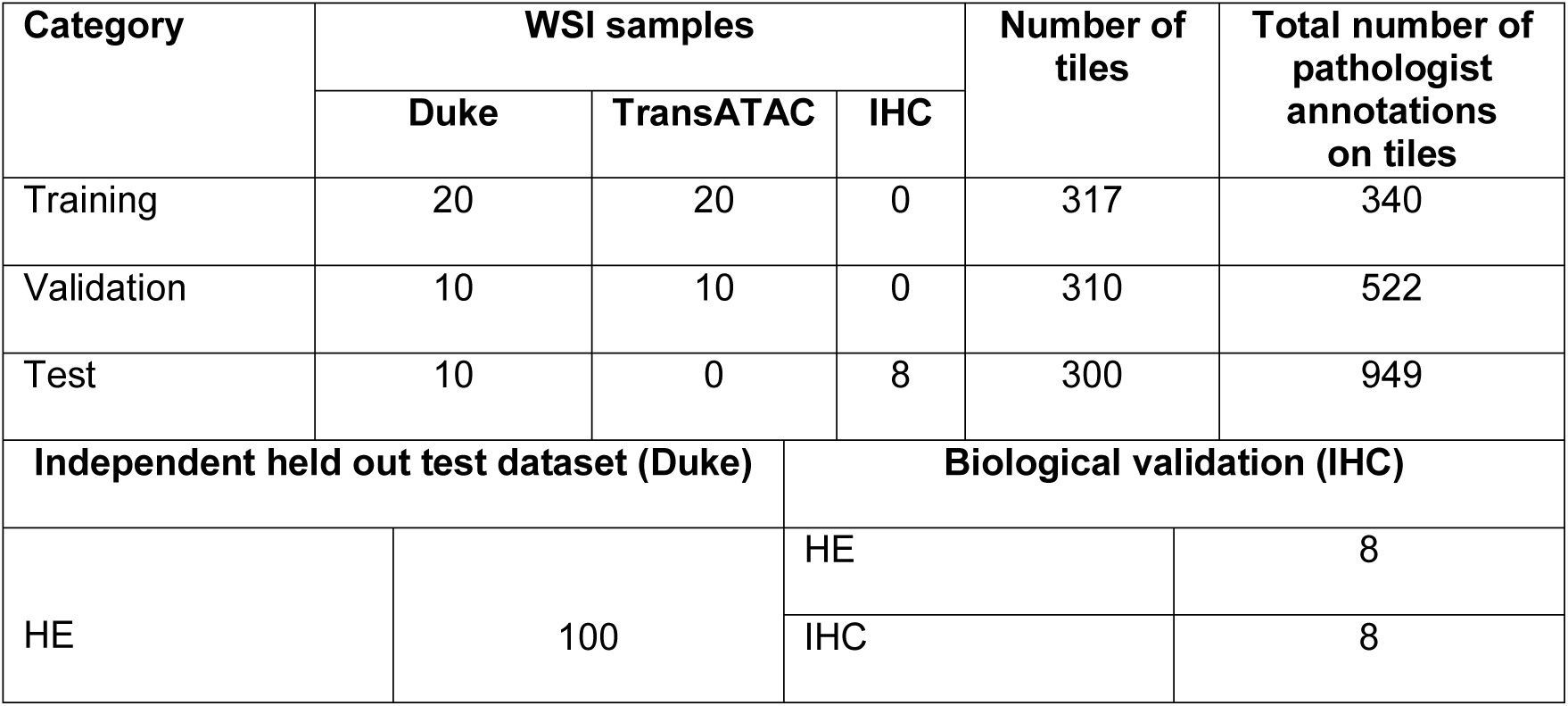
Breakdown of training and validation tiles used in annotations of DCIS.

**Table 3.**
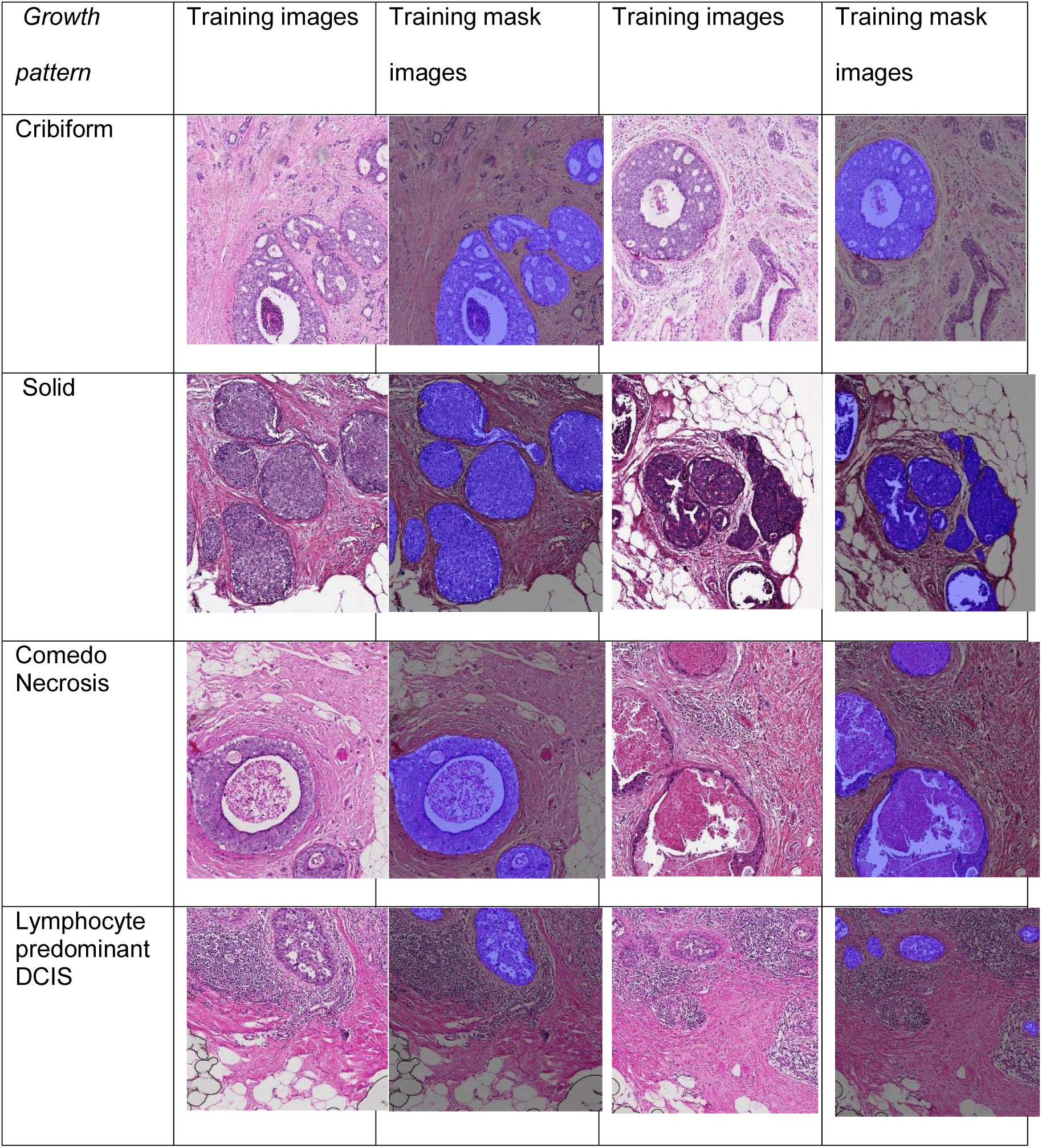
Representative examples of DCIS from the Duke and TransATAC cohorts with different growth patterns.

### UNMaSk Framework

The proposed UNMaSk (UNet-IM-Net-SCCNN) framework was designed to integrate multiple deep learning networks to: 1) identify tissue using UNet, 2) detect and segment DCIS regions using Inception MicroNet (IM-Net), 3) segment and classify single cells using MicroNet and spatially constrained convolutional neural network (SCCNN); 4) Exclusion of invasive region from DCIS adjacent images; 5) spatial tessellation to identify immune cells adjacent to individual DCIS ducts (Figure 1).

**Figure 1:**
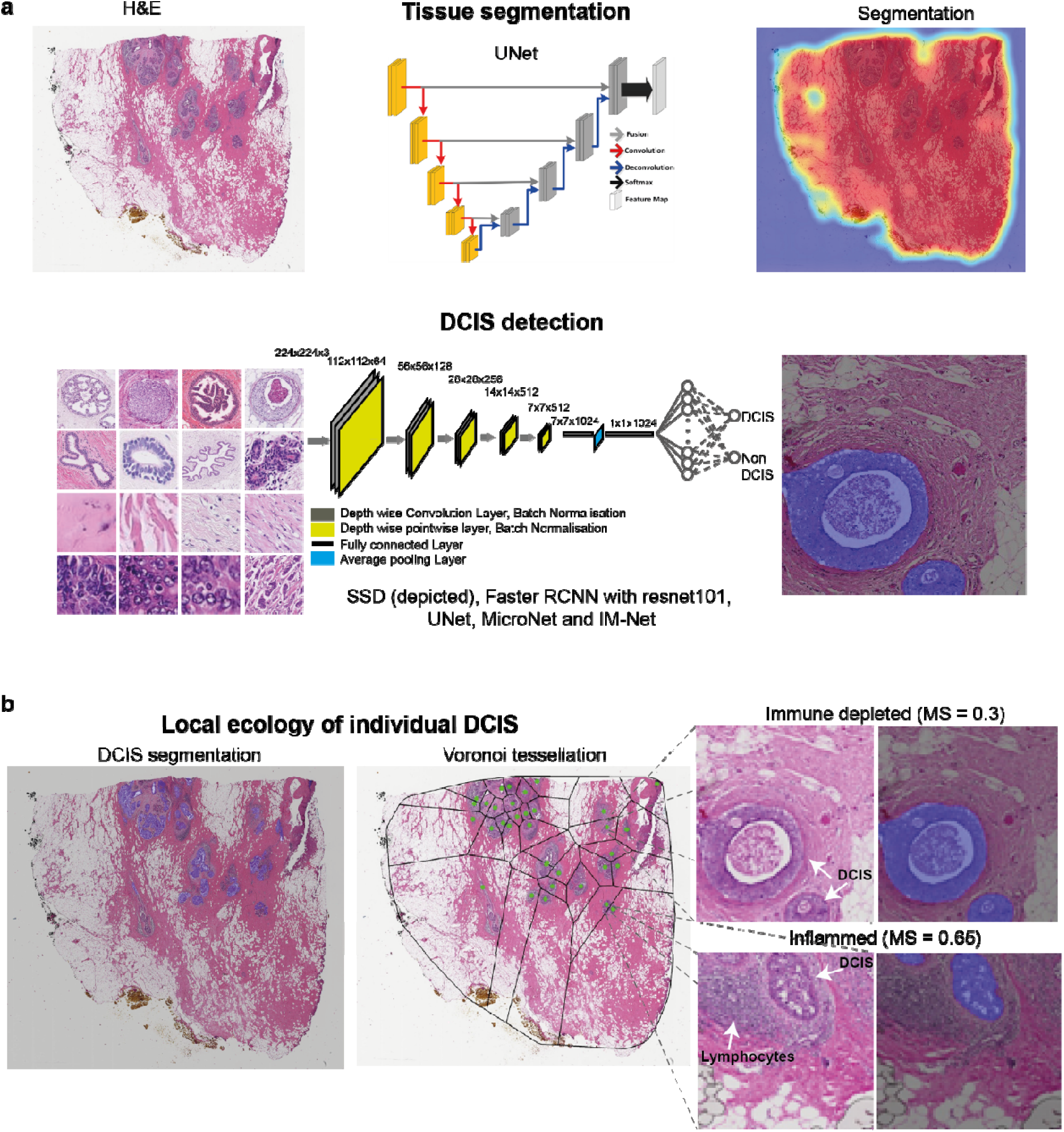
Overview of proposed UNMaSk pipeline for DCIS detection and segmentation. a. UNet architecture for tissue segmentation and one of the existing deep learning methods, single shot detector (SSD) architecture, used for DCIS detection. b. Spatial Voronoi tessellation to examine local tissue ecology for each DCIS duct, based on deep learning results on DCIS segmentation and single cell classification. Examples shown are immune predominant/inflamed and immune depleted ecology local to individual DCIS ducts from the DCIS Immune colocalisation / Morisita Score (MS) spatial analysis.

### 1. UNet for tissue detection

We used generic UNet encoder decoder architecture (18) to perform the tissue segmentation at 1.25x resolution of the image. We employed the openslide library (19) to save the images at 1.25x resolution. To fine tune the network, we performed empirical experimentation based on the patch size of 512×512. 50 training images were selected at random from the datasets, followed by standardized mean normalization. The experiments used a learning rate of 0.001 for 50 epochs with 20 percent held out dataset for the validation of the tissue segmentation. The segmentation output yielded from UNet and a threshold based segmentation indicated higher segmentation accuracy for UNet based on the DICE obtained from these approaches. Further breakdown of quantitative metrics including DICE are provided in supplementary Table S3.

### 2. IM-Net for DCIS detection

The proposed IM-Net architecture design includes contracting path which acts an encoder, and the expanding path which acts as a decoder of features from the network (Figure 2). We used five branches in the contracting path and five branches in the expanding path where data was encoded along with the spatial context with multiple inputs applied to the first three branches. To simultaneously gain local features and wider context at the same level, we introduced a custom inception block with batch normalisation performed on resized images to generate feature maps from the convolution blocks (B1, B2 and B3). Each inception block of our IM-Net (Figure 2) used the 1×1 convolution to extract features at different scales of input and the parallel paths in the inception block aid to concatenate all the features from four different scales to feed to individual blocks in the contracting path. Features from the convolution blocks were preserved and passed to the expanding path to preserve crucial low-level information. Low-level information is particularly important for segmentation and by concatenating and upsampling these features at different resolution, the network learns to preserve weak boundary features. Learning weak boundary features is a specific challenge for DCIS regions with necrosis, thus it is particularly important to address this to achieve consistent segmentation of DCIS.

**Figure 2:**
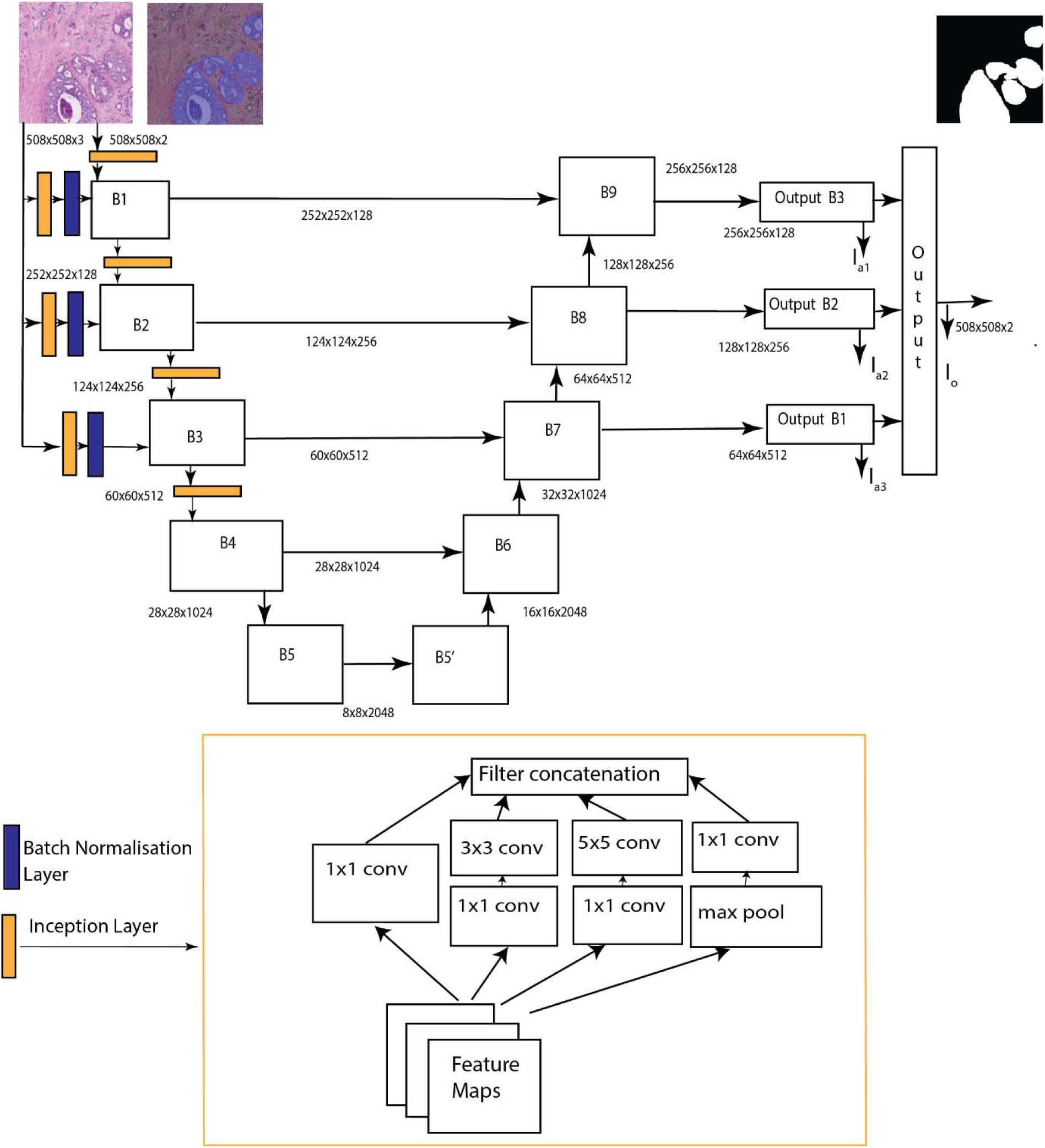
Schematic of IM-Net architecture for detection and segmentation of DCIS. Five branches in the contracting path and five branches in the expanding path encode features along with spatial context with multiple inputs applied to the first three branches. Inception blocks with batch normalisation performed on resized images generate feature maps from the convolution blocks (B1, B2 and B3). Features from the convolution blocks were preserved and passed to the expanding path to preserve crucial low-level information for DCIS boundary localisation.

Input image tiles were of size 2000×2000, from which patches of size 508×508 were extracted and augmented with random flip, rotation, scaling, gaussian blur, barrel and pincushion distortion. For the tiles at the edges of the image, we padded them with neighbouring patches to ensure all training regions were of size 508×508. They were then normalized to zero mean and unit norm and then fed to the network architecture as depicted in Figure 2. Input patches for IM-Net were set to 508×508 to capture representative context of DCIS. Selection of patches for training or validation is set to random during the training process. We used an initial learning rate (*lr*0 = 0.0001) and reduced it through the epochs (*lr = lr*0/10^*epoch*^). The loss function was defined as weighted cross entropy loss from the main and auxiliary outputs.

### 3. MicroNet and SCCNN for single cell classification

MicroNet has an encoder-decoder style architecture with multi-level input for the encoder blocks for capturing spatial context. It has been demonstrated to be capable of handling robustness in the presence of high level of noise (20). In a generic convolutional neural network framework, maxpooling blocks are often used at different stages, but this leads to information loss while aiming for semantic segmentation. To address this classical challenge, MicroNet uses receptive field and counters the loss of information by introducing an additional original image and its feature map at different input points of encoder block to guide semantic segmentation.

Single cell classification was performed using MicroNet segmentation on individual cells followed by SCCNN cell classification (8). We integrated and optimised the segmentation network to improve the detection of cells of different sizes. The centroids of individual segmented cells were estimated and then fed to SCCNN classifier. SCCNN classifier network uses neighbouring ensemble prediction combined with the standard softmax for classification. Empirical evaluation was performed on the validation dataset using random sampling of tiles across the WSI images. Furthermore, expert cell annotations were collected, and then used for evaluation of our classification on individual cells.

### 4. Exclusion of invasive region

Using the DCIS segmentations from IM-Net, we used these as regional masks, overlaid them with single cell classifications that identified epithelial cells, and reclassified epithelial cells inside DCIS regions as *in situ* epithelial cells. The rest of epithelial cells were reclassified as “other” epithelial cells that could include both invasive and normal ductal cells, and were removed from subsequent analyses. This integrative approach thus removed invasive components, leaving DCIS epithelium for subsequent spatial analysis in adjacent DCIS cases.

### 5. Spatial tessellation and Morisita index

To study the DCIS microenvironment, we used the DCIS segmentation yielded from IM-Net. In adjacent DCIS, regions comprising IDC components, were excluded after DCIS segmentation and cell classification. Furthermore, spatial compartmentalization was achieved automatically by partitioning the DCIS tissue space using Voronoi tessellation. The tessellation is a partition of space according to neighbourhood relations of a given set of points in the space. It has been suggested that Voronoi tessellation mimics the biological patterns present in the histological image and naturally emerged patterns (16). This property combined with the ecological index aids to study the ecological characteristics of individual DCIS and is similarly used in histology studies to provide spatial context of diverse cell types coexisting within the microenvironment (17).

Let *K* be a set containing all coordinates of DCIS *D* and let (*D*_*k*_)_*k*∈*K*_ be the coordinates of a DCIS*k*. A Voronoi region *R*_*k*_ generated by DCIS duct *D*_*k*_ contains all cells *P* that are not seeds and are closer to *D*_*k*_ than to any other seed *D*_*j*_, *j* ≠ *k*. Let *d*(*Q*_*i*_, *Q*_*j*_) be the Euclidean distance function between two centroids of DCIS *Q*_*i*_ and *Q*_*j*_ then

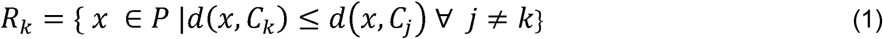

Centroids from IM-Net segmentation were estimated and the resulting binary masks were mapped back to lower resolution (1.25x) of the image. The centroids were then used as seed for calculating the Voronoi polygon. Because of these mathematical principles underlying Voronoi tessellation, lymphocytes within a polygon will be closer to its seed than any other seeds. This means that the closest DCIS duct for a lymphocyte is the one that “seeds” the polygon containing this lymphocyte. Subsequently, lymphocytes and stromal cells within *in situ* microenvironment are reclassified as *in situ* lymphocytes and *in situ* stromal cells.

Morisita-Horn similarity index is an ecological measure of community structure to quantify the extent of spatial colocalisation or overlap between two spatial variables, and we have demonstrated its use in studying cancer-immune cell colocalisation in IDCs (21). Here to measure colocalisation of DCIS and lymphocytes, the Morisita index was modified by restricting calculation in Voronoi polygons and estimating the density of epithelial cells and immune cells in the newly defined space. This space was further divided into voronoi grids, following (21). This is to provide more spatial points for calculation, and to prevent a lack of power for samples with low number of DCIS ducts. The number of immune cells and epithelial cells for each polygon *i* are denoted as 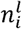 and 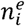, based on voronoi tessellation. Let 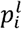 and 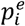 denote the fraction of immune cells and epithelial cells in polygon *i*, i.e.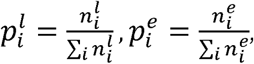, Morisita-Horn similarity index are henceforth referred as DCIS immune colocalisation score throughout the manuscript and it is calculated as:

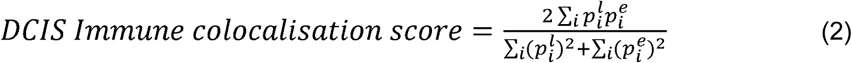

High DCIS immune colocalisation score indicates that TILs colocalise well with DCIS ducts within a sample, that is, high spatial variability; whereas, low DCIS immune colocalisation score could indicate TILs to only localise with part of the DCIS, i.e., spatial homogeneity.

### State-of-the-art networks for comparison

Single shot detector (22), resnet 101 based RFCNN network (23), UNet (18) and MicroNet (20) were trained following optimal configuration as described in the original publications. Network parameters for input patch size varied from 224 to 508 following original implementation. Hyperparameters such as learning rate varied from 0.00001 to 0.001 based on both original publications and experiments. Further details on number of parameters, optimizer and learning rate for individual network of choice are provided in supplementary Table S1.

### Biological validation

CK5 is an immunohistochemistry myoepithelial marker routinely used in the clinic due to its high sensitivity in the detection of DCIS (24). To further evaluate the DCIS segmentation accuracy on H&E images, we cut whole-tumour serial sections from 8 samples and stained them, resulting in 8 pairs of H&E and CK5 immunohistochemistry slides (Dako, catalogue number M7237, lot number 20014544). Based on morphology and localisation supported by CK5 expression, areas of DCIS were hand annotated, providing quantity and area of individual DCIS for a quantitative evaluation based on correlation of the automated segmentation methods on H&E images.

### Statistical methods

Wilcoxon test was used to determine statistical significance of differences in immune scores between pure and adjacent DCIS samples, and p-value < 0.05 was used to determine significance. To further test the difference at patient level, one WSI for a patient were sampled to represent a patient, and randomized over 100 times iteratively. Then the number of times Wilcoxon test reached significance were calculated. Correlation analyses of parameters such as estimated number of DCIS regions and estimated area of DCIS was determined between automated H&E method and hand annotations by IHC marker. All correlation tests used Spearman’s correlation method. All statistical tests were carried out in R.

### Quantitative evaluation at slide level

We used DICE coefficient for our quantitative evaluation on the test dataset with the ground truth annotation. DICE measures the spatial overlap between the ground truth annotation and automated segmentation methods. Quantitative measures include DICE, positive predictive value (PPV), negative predictive value (NPV), true positive rate (TPR), true negative rate (TNR), false positive rate (FPR) and false negative rate (FNR) for each test dataset across all slides.

## Results

### IM-Net for DCIS detection and segmentation

To automate the identification and segmentation of morphologically heterogeneous DCIS ducts in H&E, we specifically designed a new deep learning framework, IM-Net, to: 1) distinguish DCIS from other tissue components by combining high level spatial context and local features using multiple inputs to the encoders, 2) provide precision in localising DCIS boundary by learning weak boundary features through concatenating and upsampling of features across spatial resolutions, 3) reduce sensitivity to tissue artefacts and local noise by using multiple filters in inception blocks. We first compared the performance of the proposed IM-Net in DCIS detection and segmentation with four state-of-the-art deep learning methods using images generated across three datasets (Figure 1a-b). These include some of the most widely used methods for segmentation or detection: single shot detector (22), resnet 101 based RFCNN network (23), UNet (18), MicroNet (20) and the proposed IM-Net. For all DCIS detection and segmentation pipelines, UNet was first used for tissue segmentation before DCIS detection (Figure 1a). Independent samples were used for training, validation, and testing, consisting of 340, 522 and 949 annotations on 40, 20, and 18 H&E whole-section images, respectively (Table 2). Three experiments were conducted to test: 1) accuracy of DCIS detection as a binary classification problem; 2) accuracy of DCIS segmentation by quantifying the overlap between pathologists’ delineation and automated segmentation; 3) validation using immunohistochemistry.

Whilst IM-Net, MicroNet, UNet and Faster RCNN yielded higher precision compared to SSD in the testing set, IM-Net achieved the highest recall (0.75, Table 4). This suggests that IM-Net can detect the highest proportion of DCIS annotated by pathologists, with potentially low false positive rates. This is confirmed by further breakdown of the false positive and true positive rates (Supplementary Table S2), which showed the lowest false positive rate as well as highest true positive rate for IM-Net among all networks (0.10 and 0.77, respectively). Secondly, the overlap between pathologists’ annotated DCIS and automated segmentation results indicated that IM-Net followed by MicroNet achieved the highest DICE (Figure 3, Supplementary Table S2). Spatial overlap estimated by DICE for IM-Net was 0.83, where DICE of 0.6 and above is typically considered to indicate a good agreement between ground truth and prediction (25).

**Table 4:** Quantitative comparison with state-of-the-art methods and the classification accuracy of deep learning networks for DCIS detection on validation dataset.

| **Model** | **Precision** | **Recall** | **F1-score** |
| --- | --- | --- | --- |
| SSD | 0.79 | 0.67 | 0.73 |
| Faster RCNN with Resnet | 0.81 | 0.65 | 0.74 |
| UNet | 0.79 | 0.64 | 0.70 |
| MicroNet | 0.82 | 0.72 | 0.75 |
| IM-Net | 0.83 | 0.75 | 0.76 |

**Figure 3:**
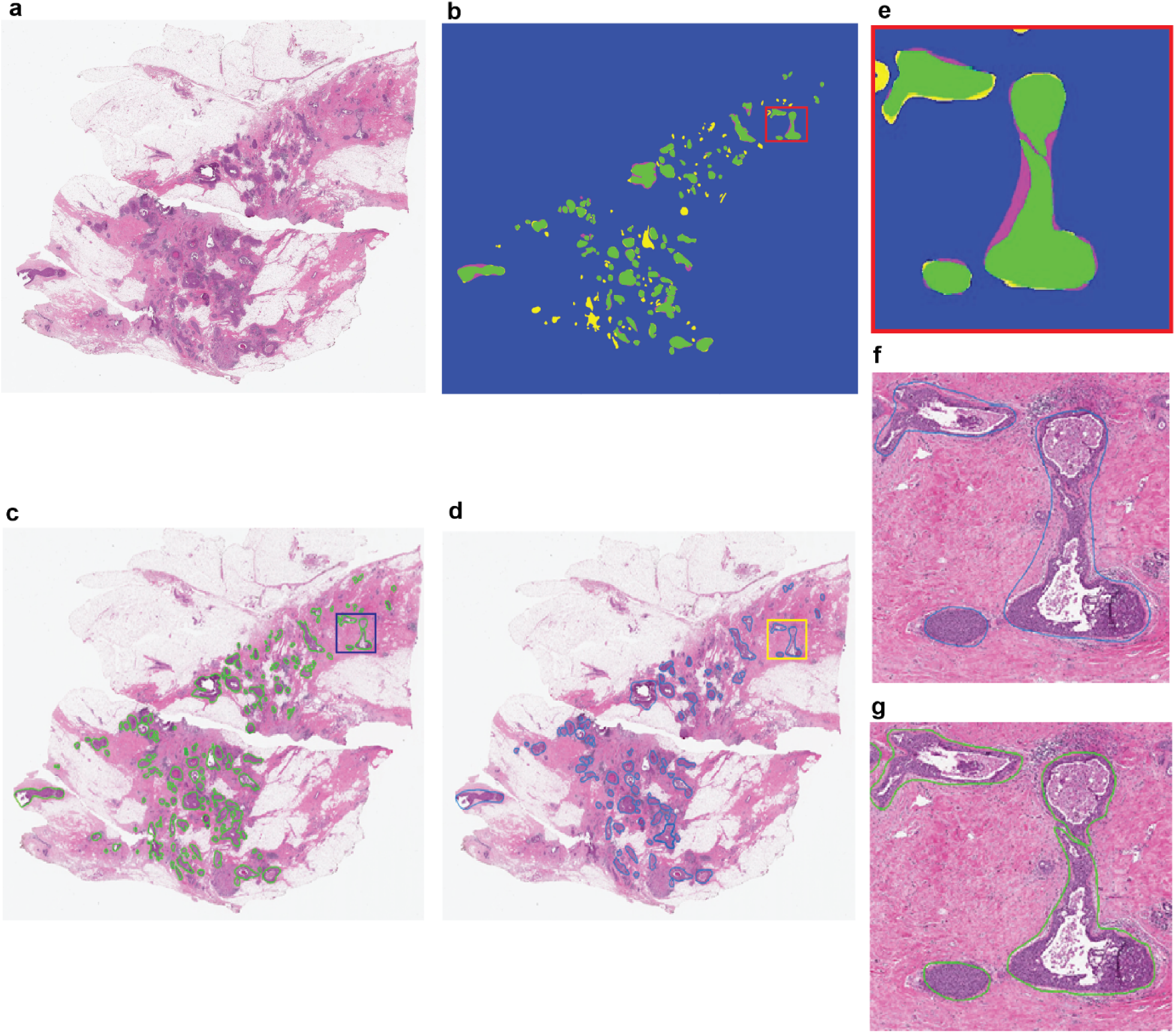
Representative H&E image with DCIS segmentation output from IM-Net.(a) H&E image. (b) Spatial overlap between the pathologist annotation and DCIS detection using IM-Net were estimated using Dice coefficient and its respective representation. True positive (TP) in green where the expert annotations match the detected area; False Negative (FN) in magenta that cover DCIS regions in annotation and not covered by DCIS detection. False positive (FP) in yellow pixels falsely segmented as DCIS and not in expert annotation. True Negative (TN) represented in blue are pixels correctly detected as background that is tissue background in both expert and DCIS detection. Inspection of the false positive regions showed that some of these were real DCIS but contained tissue artefacts or tears, which was the reason why pathologist did not annotate them. (c) DCIS segmentation based on IM-Net. (d) Pathologist annotation of DCIS on H&E image. (e) Region of interest depicting the Dice image map. (f and g) Region of interest depicted from pathologist and IM-Net approach respectively.

Further examination of the false positives of DCIS detected by IM-Net indicated that a number of these were due to tissue artefacts or tears occurring within or near DCIS, which prevented pathologists from annotating them. However, these were still detected by automated methods, and upon consultation with pathologist’s we arrived at an agreement on these difficult cases. In these few images, we observed highly clustered DCIS regions and tissue artefacts that led to the difficulty in annotating all DCIS regions in the WSI. We therefore performed immunohistochemistry experiments to base our validation on biological marker-based expression (Methods). This biological validation demonstrated a high correlation between IM-Net segmentation of DCIS with hand annotation based on cells positive for CK5 expression lining the basal membrane (cor=0.99 vs cor=0.98-0.94 for other networks, Table 5). Superior performance of IM-Net was further supported in quantitative analysis of the overlap of detected and annotated DCIS areas (DICE = 0.83 for IM-Net; DICE = 0.82 for MicroNet vs DICE = 0.65 – 0.77 for other networks (Supplementary Table S1). Visual inspection confirmed the quantitative data, showing a good agreement between automated detection on the H&E and CK5-based annotations on serial IHC, irrespective of different growth patterns (Figure 4).

**Table 5.** Correlation between number of DCIS and area of DCIS estimated from biological ground truth.

|  | **IM-Net** | MicroNet | UNet | RFCNN | SSD |
| --- | --- | --- | --- | --- | --- |
| Number of DCIS | **0.97** | 0.97 | 0.95 | 0.93 | 0.92 |
| Estimated Area | **0.99** | 0.99 | 0.98 | 0.96 | 0.94 |

**Figure 4:**
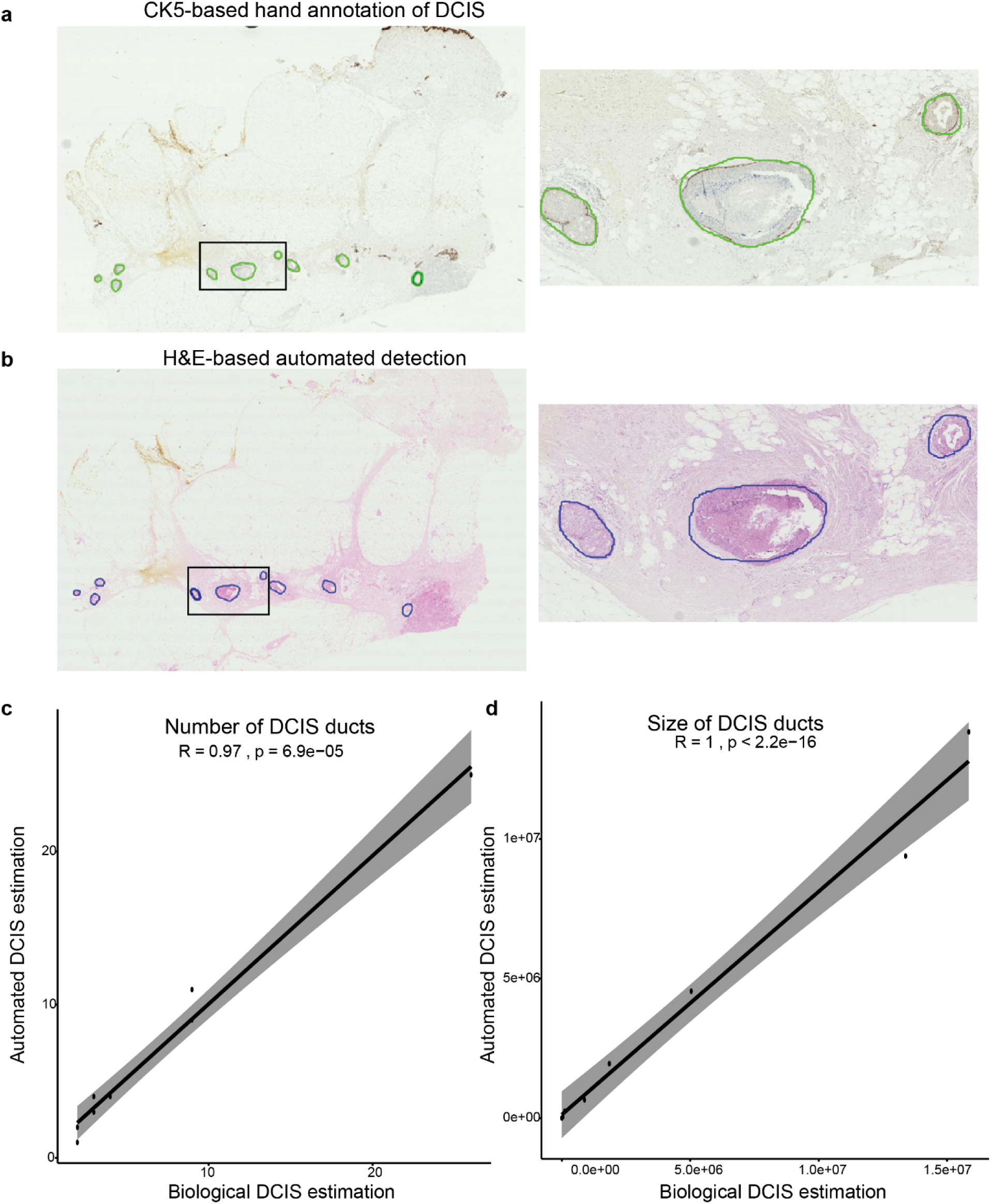
Biological validation of automated DCIS detection using CK5 immunohistochemistry. (a) An example showing CK5 IHC image where DCIS regions were annotated by hand following the CK5 expression pattern, indicated by green contour. (b) Segmented H&E image with DCIS regions marked in blue contour by IM-Net for the same sample. (c) Quantitative assessment of the IHC-H&E correlation using H&E-based automated DCIS detection result and hand annotations on IHC using estimated number of DCIS. (d) Quantitative assessment of the IHC-H&E correlation using H&E-based automated DCIS detection result and hand annotations on IHC using estimated area of DCIS.

### TILs distribution mapped by deep learning

To identify immune cells and stromal cells that form the immediate microenvironment of DCIS, we used an established deep learning approach for identifying and classifying single cells. It uses an established MicroNet approach for cell segmentation by localising the centre of an individual cell in the H&E image tiles and then classifying the cell based on our implementation of SCCNN classification network. For training the classifier of individual cells, we used 12 WSI and tested on 3 WSI comprising 11,412 annotations for cells during training and 5,207 cells for validation. Accuracy of classification was evaluated by 3-fold cross validation on the training set of individual cells where we obtained 89.3% accuracy, we obtained an accuracy of 87.1% on an independent validation dataset. As a result, all the cells were mapped in the H&E images and classified into epithelial, fibroblasts, and lymphocytes (Figure 5a-c).

**Figure 5.**
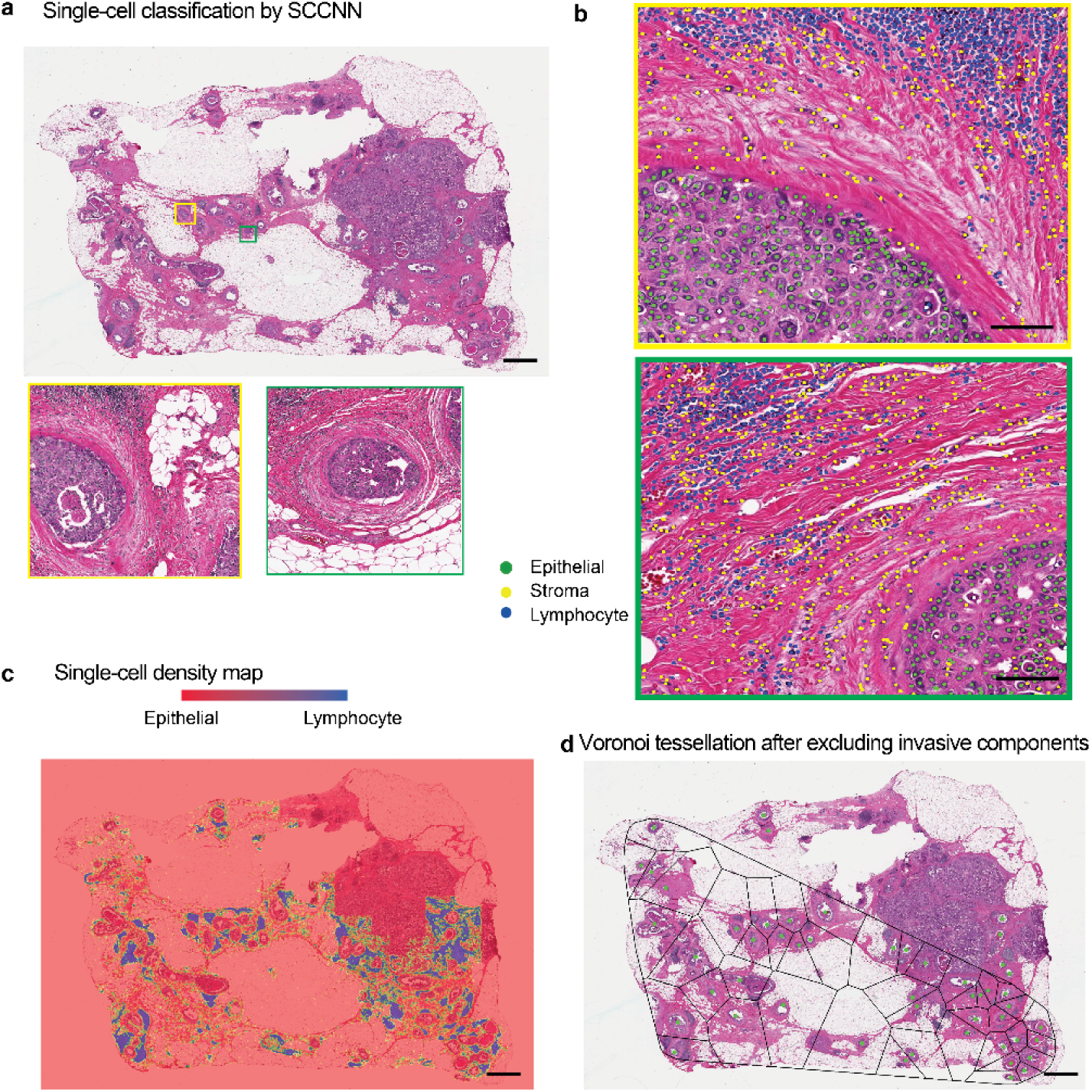
UnMaSk with single cell classification excluding invasive region single cell classification using DCIS segmentation and voronoi tessellation. (a) A representative example of an adjacent DCIS case illustrating single-cell classification results in two DCIS regions. Scale bar represents 100⎧m. (b) High resolution images of areas within the two DCIS regions, showing single cell classification using unified segmentation and classification pipeline based on MicroNet and SCCNN, classifying cells into epithelial cell (green), stromal cell (yellow) and lymphocyte (blue). Scale bar represents 50⎧m. (c) Heatmap showing lymphocyte cell density based on single-cell classification results. (d) Voronoi tessellation using the centres of DCIS ducts as seeds, performed over tissue space excluding epithelial cells identified by single-cell classification that were not DCIS. Because of the mathematical principles underlying Voronoi tessellations, lymphocytes within a polygon will be closer to its seed than any other seeds. This means that each lymphocyte can now be assigned to its closest DCIS duct within the tessellation space, thereby quantifying lymphocyte abundance for each DCIS duct locally. Note that because convex polygon was used, some of the DCIS ducts closer to invasive region were omitted from the analysis. Scale bar represents 100µm.

### Increased co-localisation of TILs and DCIS in IDC

Spatial variability of TILs among DCIS ducts could inform differential ecological features that ultimately dictate invasive potential and fundamental biological underpinning. However, this would require quantitative methods to examine the spatial heterogeneity, instead of focusing on only abundance of lymphocyte cells. Therefore, to characterise local tumour ecology of individual DCIS, we then applied a spatial statistical method of colocalisation, the DCIS immune colocalisation score, previously shown to be prognostic for Her2-positive breast cancer (21) using known cellular phenotypes. Widely applied in ecology, this spatial statistical method is often used to examine spatial overlap/colocalisation, providing a single score for a spatial area: a high score indicates that the two spatial variables tend to co-localise, whereas a low score suggest spatial separation. We applied this method to the Duke dataset excluding training/validation samples and samples with less than 5 DCIS ducts, to compare co-localisation pattern of TILs and between pure DCIS samples and DCIS adjacent to IDCs in a total of 92 WSIs, representing n = 26 pure and n = 22 adjacent DCIS cases (Figure 6a-f).

**Figure 6.**
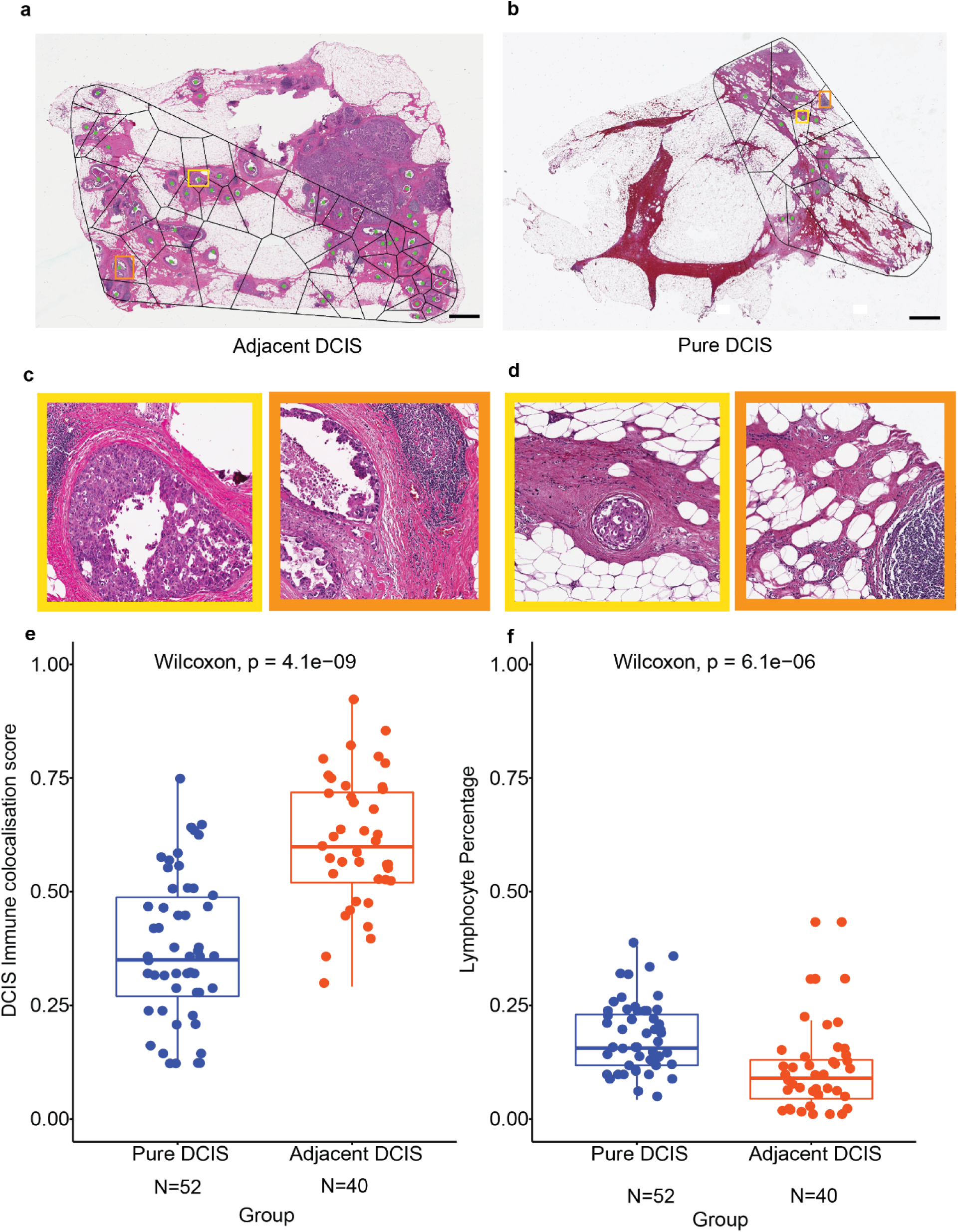
Comparison of TIL distribution pattern local to DCIS ducts in adjacent versus pure DCIS cases. (a) Voronoi tessellation of adjacent DCIS excluding invasive components (b) Voronoi tessellation of Pure DCIS. Scale bar represents 100⎧m. (c) Representative DCIS region enclosed within the voronoi of adjacent DCIS. (d) Representative DCIS region enclosed within the voronoi of Pure DCIS. (e) Boxplot illustrating the difference in DCIS immune colocalisation score calculated using the Morisita index. It was computed by associating individual DCIS duct with the surrounding lymphocyte within the voronoi region; a high score indicates the spatial colocalisation of lymphocytes and DCIS ducts. Each point corresponds to a WSI image, 52 WSI from n=40 patients in the pure DCIS and 40 WSI from n=25 patients in the adjacent DCIS group. (f) Box plot illustrating the difference in overall lymphocyte percentage in all cells for WSIs of pure DCIS and adjacent DCIS cases (after exclusion of invasive tumour regions), using only single-cell classifications.

Despite significantly higher numbers of TILs (measured as percentage of lymphocytes in all cells after exclusion of invasive tumour regions if any) in pure DCIS subjects than in adjacent DCIS (p = 6.1e-06), the DCIS immune colocalisation score showed that co-localisation of TILs was lower in these patients compared with samples from adjacent DCIS from patients with IDC (p = 4.1e-09, Figure 6e-f). The difference in DCIS immune colocalisation score remained significant when the DCIS immune colocalisation score were averaged per patient (p = 2.7e-06, Figure S1a), randomly sampled from for cases with more than 1 slides available (p < 0.05 for 100 of 100 random sampling), or randomly sampling from pure cases to match the number of adjacent cases (Figure S1b). This suggests that although there were more TILs in the pure DCIS samples, these TILs did not co-localise well with DCIS ducts. In contrast, TILs in DCIS sampled from tissues adjacent to IDCs tend to localise with DCIS, possibly due to heightened inflammatory response. Representative examples with positioning of lymphoid aggregates in pure DCIS and adjacent DCIS samples are shown in Figure 6c-d. There was no correlation between TILs abundance and the DCIS immune colocalisation score within each DCIS group (p = 0.082 for pure DCIS and p = 0.21 for adjacent DCIS), thus the spatial pattern was not dependent on TIL abundance. These quantitative data and empirical observations support the different types of lymphocyte/epithelial interactions in these two types of DCIS, and further highlights the importance of examining spatial heterogeneity beyond cellular abundance alone.

Furthermore, we tested the dependency of the DCIS immune colocalisation score on clinical parameters of these patients including ER, PR, HER2, age and grade (Table 1). Firstly, the DCIS immune colocalisation score continued to differentiate two DCIS types within ER-positive and ER-negative patients (Figure S2). Secondly, in a multivariate model predicting DCIS types with the DCIS immune colocalisation score, TILs abundance, and clinical parameters, the DCIS immune colocalisation score remained a significant discriminator of pure, versus adjacent DCIS (p=9.08e-03). These data suggest that the DCIS immune colocalisation score which measures the colocalisation pattern of TILs and DCIS ducts within the tissue, may have utility in differentiating pure and adjacent DCIS, independent of known clinical parameters.

## Discussion

There is a high variability in clinical outcomes and propensity for invasion among DCIS cases (1-3). Development of reliable markers which can identify DCIS patients who are likely to follow a benign course from those who would benefit from therapy are currently an unmet requirement for clinical care. To provide advanced tools for identifying potential interactions between individual DCIS and its microenvironment, we developed a deep learning pipeline, UNMaSk, that integrates tissue segmentation, DCIS segmentation, single cell classification and spatial analysis in routine H&E histology images. An innovative use of this pipeline, for the study of tissue inflammatory microecology using a spatial statistical measure, the DCIS immune colocalisation score, quantifies the spatial colocalisation of individual DCIS ducts and immune cells. As such, our approach can be used to deliver a DCIS immune colocalisation score that integrates morphological context of DCIS and high-resolution single cell based classification with intrinsically captured heterogeneous DCIS microenvironment. This type of integrative approaches was not previously possible due to the lack of computational utility, and we bridged this gap in this study.

Routine clinical H&E-stained images are known to be highly noisy with staining variability and preprocessing artefacts. Frequently occurring artefacts in WSI are fixation problems, coverslip artefact and pen marking at the margin of the fatty tissue that represent real world diagnostic breast histology images. Our framework incorporates problem-specific design using tissue segmentation based on UNet to alleviate effects due to artefacts. Training and evaluation were performed on cohorts that were processed, stained and scanned in independent laboratories using different digital slide scanners, to reduce overfitting and improve generalizability across scanner and datasets in an unbiased and systematic manner. A potential utility of UNMaSk is therefore detection of DCIS as part of a screening program, which warrants further investigations.

Additionally, we showed that the proposed network (IM-Net) for DCIS segmentation was able to capture and better generalize image semantics across granularities with the usage of inception block and the multiple resized input images fed to each convolution block in the contracting path. The tailored inception block usage in the contracting path and its effective use of the weighted loss function introduced for the first three convolution and transpose convolution blocks preserved features from DCIS. Despite the major challenges in analysing diverse growth patterns across images, the proposed IM-Net has improved DICE and reduced false positive rate compared to the other networks (Supplementary Table S1). The context features for large and small DCIS were captured from multiple resized images and the IM-Net was found to be agnostic to the size and morphology of DCIS. Furthermore, IM-Net was capable of capturing weak boundaries of the duct containing necrosis without over-segmentation of DCIS regions.

The use of a validated ecological index combined with the morphological patterns of DCIS yielded from the UNMaSk pipeline demonstrates a potential innovative use of this tool. The ability to derive ecological parameters such as lymphocytic colocalisation with high spatial resolution regions comprising only DCIS was not possible without a fully automated framework like UNMaSk. Spatial mapping of individual DCIS ducts combined with single-cell classification based on deep learning enabled the characterization of DCIS tissue habitat at micro scale, i.e. microecology. By excluding the invasive regions in the adjacent DCIS cases, the proposed UNMaSk focuses on DCIS components and extrapolate statistical inference to predict lymphocyte association with individual ducts in both pure and adjacent DCIS cases.

In the Duke cohort, we observed that there is a lower level of overall TILs but higher colocalisation of TILs and DCIS ducts in DCIS adjacent to invasive cancer compared to pure DCIS cases. Increased number of leukocytes is often associated with high grade DCIS, Her2 status, and IDC (11). Our data showing lower TIL abundance in adjacent DCIS compared with pure DCIS may be attributed to the fact that pure DCIS samples in our cohort were intermediate to high grade (47% grade 2 and 53% grade 3), and that invasive tumour regions were excluded in the calculation for the adjacent cases. Nevertheless, our data showing differential colocalisation spatial pattern is new. Although high spatial variability of TIL distribution in DCIS has been reported, it has rarely been quantitatively measured. Our data on an increased level of spatial colocalisation of TILs to individual DCIS ducts in adjacent DCIS suggest a different immune microenvironment in these patients compared with pure DCIS, and that an increase in the colocalisation spatial immune score may predict which DCIS is likely to progress to invasive disease. This will require further validation in larger cohorts, as well as in samples with longitudinal follow up. Nevertheless, this data is consistent with a study where dense chronic inflammation surrounding DCIS, defined as circumferential cuffing of the duct by lymphocytes or plasma cells at least three cell layers in thickness, was associated with a high recurrence score of Oncotype DX DCIS (26). This preliminary evidence supports future studies to validate the use of the DCIS immune colocalisation score in predicting propensity of individual DCIS for invasive progression. This hypothesis is biologically intriguing, and these studies are currently under way in our lab.

Future opportunities include the generalization of our deep learning method to other types of histological images such as immunohistochemistry for defining immune cell subsets surrounding DCIS, investigation of the morphological and architecture details within detected DCIS ducts to account for the types of DCIS, integration with genomic aberrations that could drive progression (17, 27), and scaling up in large patient cohort analyses. Thus, our study contributes to driving future research towards the use of novel parameters such as spatial immune score based on automated histology image analysis for the identification of drivers and biomarkers of progression from DCIS to invasive cancers. Additional studies using these approaches with matched follow-up cases may be useful for validation and to assess the relevance of DCIS immune colocalisation score with respect to the recurrence. This could further potentially aid in stratifying risk of progression, and ultimately improve personalised clinical care.

## Conclusions

We presented a new deep learning pipeline, UNMaSk, for the automated detection and segmentation of DCIS ducts. Our comprehensive evaluation experiments on three sample cohorts, using expert annotations and biological immunohistochemistry, and comparison with state-of-the-art convolutional networks demonstrated UNMaSk to be agnostic to diverse size and growth patterns of DCIS. Our framework allowed the integration of spatially heterogeneous DCIS to be associated with lymphocyte distribution by considering topological arrangement of DCIS. This provides a unique opportunity to study inflammatory response reactive to carcinoma at high spatial resolution, paving the way for quantitative analysis of DCIS ecology and morphology, and AI-aided risk stratification for DCIS disease.

## Data and code availability

All training data, including the fully anonymised raw H&E image tiles and pathological annotations as binary marks, as well as Python code, will be made available upon publication.

## Acknowledgments

This project was funded by Breast Cancer Now (2015NovPR638), NIH R01 CA185138 and CDMRP Breast Cancer Research Program Award BC132057. Y.Y. also acknowledges funding from Cancer Research UK Career Establishment Award (C45982/A21808), Children’s Cancer and Leukaemia Group (CCLGA201906), NIH U54 CA217376, CRUK Brain Cancer Award (TARGET-GBM), European Commission ITN (H2020-MSCA-ITN-2019), Wellcome Trust (105104/Z/14/Z), and The Royal Marsden/ICR National Institute of Health Research Biomedical Research Centre. C.C.M. was supported in part by NIH grants U54 CA217376, U2C CA233254, P01 CA91955, R01 CA170595, R01 CA185138 and R01 CA140657 as well as CDMRP Breast Cancer Research Program Award BC132057 and the Arizona Biomedical Research Commission grant ADHS18-198847. We thank the ATAC/LATTE trailists and we thank Andrew Dodson for technical support. ESH was supported in part by NIH grants U2C CA17035, R01 CA170595, R01 CA185138 as well as CDMRP Breast Cancer Research Program Award BC132057 and the CRUK Grand Challenge Award.

