## Supplementary Tables and Figures for "Unmasking the tissue microecology of ductal carcinoma *in situ* with deep learning"

**Table S1.** Details of patch size and hyper-parameters for DCIS detection and segmentation.

| Model | Input patch size | Total number of parameters | Learning Rate | optimizer |
| --- | --- | --- | --- | --- |
| SSD | 224x224 | 27,304,700 | 0.0001 | RMSprop optimizer |
| RFCNN | 224x224 | 68,607,459 | 0.0003 | Momentum optimizer |
| UNet | 252x252 | 15,394,825 | 0.001 | Adagrad optimizer |
| MicroNet | 508x508 | 912,514,125 | 0.001 | Adagrad optimizer |
| IM-Net | 508x508 | 913,415,373 | 0.0001 | Adagrad optimizer |

**Table S2.** Breakdown of detection and segmentation performance of convolution networks on 18 whole slide images indicating accuracy.

| **Model** | **TPR** | **TNR** | **FNR** | **FPR** | **PPV** | **NPV** | **Dice** |
| --- | --- | --- | --- | --- | --- | --- | --- |
| SSD | 0.70 *±* 0.17 | 0.95 *±* 0.09 | 0.24 *±* 0.05 | 0.10 *±* 0.15 | 69.29 *±* 0.2 | 94.59 *±* 0.2 | 0.65 *±* 0.7 |
| Faster RCNN with ResNet | 0.71 *±* 0.16 | 0.98 *±* 0.04 | 0.27 *±* 0.06 | 0.04 *±* 0.02 | 71.2 *±* 0.15 | 97.75 *±* 0.03 | 0.71 *±* 0.65 |
| UNet | 0.73 *±* 0.18 | 0.97 *±* 0.08 | 0.28 *±* 0.05 | 0.08 *±* 0.10 | 72.8 *±* 0.20 | 98.2 *±* 0.07 | 0.77 *±* 0.30 |
| MicroNet | 0.75 *±* 0.12 | 0.98 *±* 0.02 | 0.24 *±* 0.03 | 0.013 *±* 0.01 | 75.79 *±* 0.3 | 98.60 *±* 0.03 | 0.82 *±* 0.50 |
| **IM-Net** | **0.77 *±* 0.10** | **0.98 *±* 0.05** | **0.24 *±* 0.01** | **0.011 *±* 0.02** | **76.8 *±* 0.1** | **98.50 *±* 0.01** | **0.83 *±* 0.30** |

**Table S3.** Breakdown of segmentation performance of tissue segmentation indicating segmentation accuracy based on Dice coefficient

| **Model** | **TPR** | **TNR** | **FNR** | **FPR** | **PPV** | **NPV** | **Dice** |
| --- | --- | --- | --- | --- | --- | --- | --- |
| Threshold | 0.72*±*0.10 | 0.50*±*0.10 | 0.27*±*0.10 | 0.49*±*0.10 | 72.4*±*10.2 | 50.7*±*10.8 | 0.73*±*0.15 |
| **UNet** | 0.72*±*0.10 | 0.76*±*0.06 | 0.28*±*0.10 | **0.23*±*0.06** | **71.8*±*10.5** | **76.5*±*6.06** | **0.78*±*0.10** |

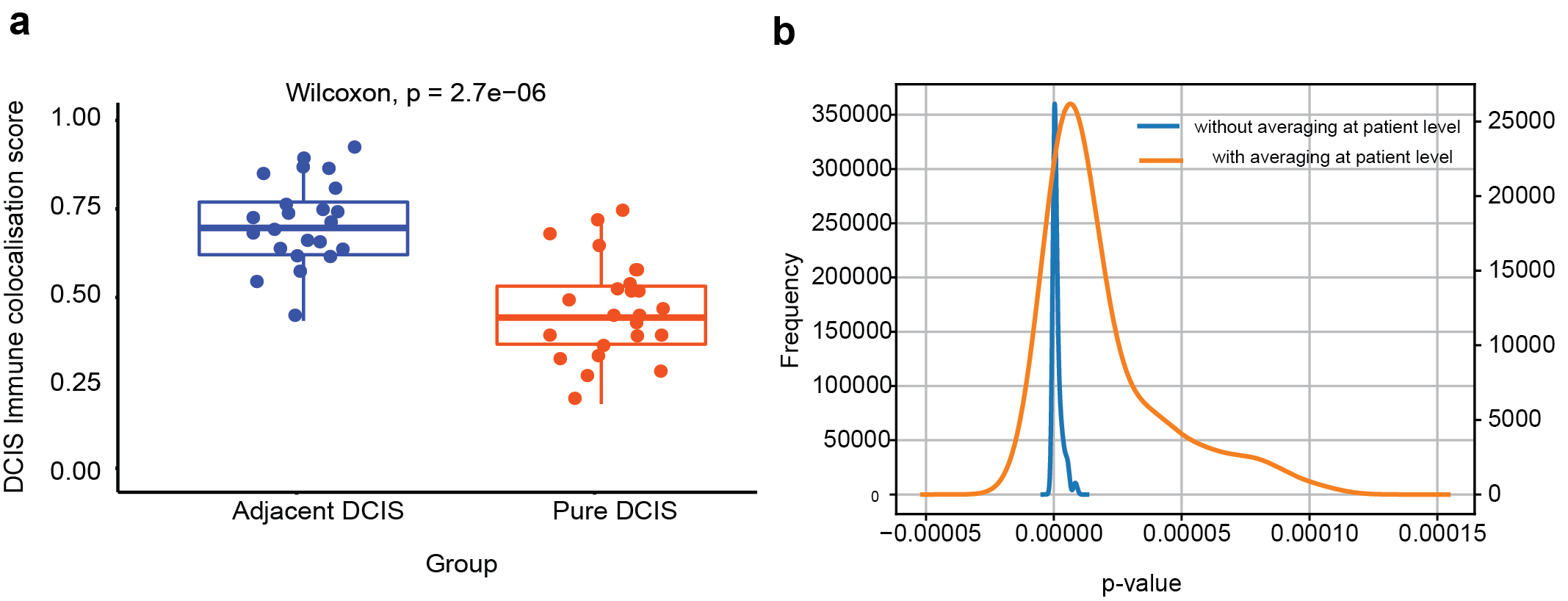

**Figure S1.** Comparison of TIL distribution pattern local to DCIS ducts in adjacent versus pure

DCIS cases after averaging per patient or by means of random sampling per patient. (a) Patient

level Morisita score was averaged from one/two/three images per patient belonging to 40 patients of

pure DCIS and 25 patients of adjacent DCIS in total. (b) Distribution of p-values in 100 random

sampling for calculated patient level scores.

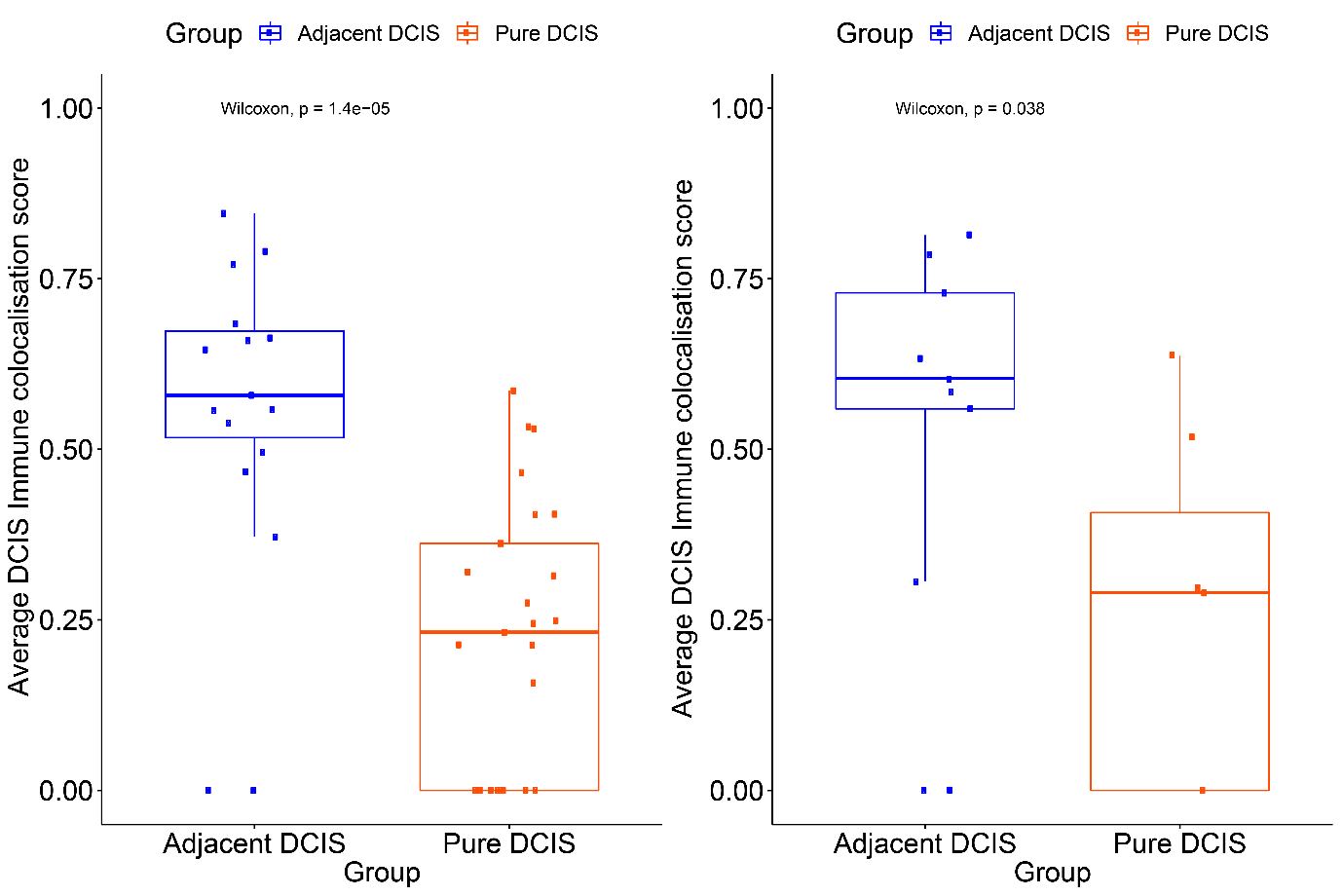

**Figure S2.** Boxplots illustrating the average difference in DCIS-immune colocalisation score at patient level between pure DCIS and adjacent DCIS for ER+ (left) and ER- (right) tumours.
